# Caste-specific ageing emerges from the evolution of resource allocation in eusocial insects

**DOI:** 10.64898/2026.06.16.732452

**Authors:** Jan J. Kreider, Thijs Janzen, Boris H. Kramer, Ido Pen

## Abstract

Eusocial insects have extreme intraspecific lifespan variation, where queens are long-lived (up to 30 years) whereas workers only live for a few months or years at most. Several studies have invoked the disposable soma theory to explain the evolution of caste-specific ageing in eusocial insects, which proposes that senescence results from a resource allocation trade-off between maintenance vs. reproduction. An extension of this theory to eusocial insects is that caste-specific ageing could emerge from a resource allocation trade-off between castes. However, to date this idea has not been formalised in a theoretical model. Here, we present an individual-based model for the evolution of ageing in social insects. In our model, queens and workers die when their nutritional state becomes too low. The evolving trait in our model is the age-specific resource allocation of individual workers, who can allocate resources between themselves, other workers, and the queen. We find that lifespan differences between queens and workers emerge from the evolved resource allocation within colonies, which are within the range of empirically observed lifespans of queens and workers in monogynous eusocial insects. Caste-specific ageing evolves in our model because queens obtain large amounts of resources, which allows them to be long-lived and highly fertile, whereas workers evolve to give resources away to enhance the queen’s reproduction and thereby their own indirect fitness. We also observe that age polyethism emerges, where young workers nurse the brood and older workers forage. Overall, our model demonstrates that both caste-specific ageing and age-related worker division of labour emerge as a consequence of evolved within-colony resource allocation.

## Introduction

Eusocial insects typically have one reproductive queen which lays all eggs while she is supported by many sterile workers. The lifespans of queens and workers differ remarkably. For instance, in some species of ants, queens become up to 30 years old whereas the workers only live for a few months or years (Keller, 1998; Kramer & Schaible, 2013a).

The evolution of caste-specific ageing has long been assumed to result from differential exposure of queens and workers to extrinsic mortality risk (Keller & Genoud, 1997; Jaimes-Nino & Oettler, 2025). Queens are usually sheltered in the nest whereas the workers are exposed to high mortality risks during foraging and defending the nest. The classical theory for the evolution of ageing (Medawar, 1952; Williams, 1957; Hamilton, 1966) predicts that such differential exposure to extrinsic mortality risk should cause the evolution of differences in the onset of senescence. This is because exposure to extrinsic mortality risk weakens selection against mutations affecting survival in old individuals (Moorad & Promislow, 2010; Gaillard & Lemaître, 2017; De Vries *et al*., 2023). Evolutionary models have challenged this verbal explanation, demonstrating that caste-specific ageing evolves because selection operates at different strengths on queen and worker survival due to reproductive division of labour (Kramer *et al*., 2022). Since the queen produces the offspring and her workers raise them, the reproductive success of a social insect colony depends more strongly on the survival of the queen than on the survival of an age cohort of workers. Therefore, selection against deleterious survival-reducing mutations is stronger in queens than in workers, which paves the way for to the evolution of long-lived queens and shorter-lived workers (Kramer *et al*., 2022). These differences in the strength of selection between castes also explain why between-caste pleiotropy, e.g. genetic correlation between queen and worker survival, reduces worker survival more strongly than queen survival (Kreider *et al*., 2021). Such genetic correlations can be interpreted as causing a between-caste trade-off, which suggests that between-caste resource allocation trade-offs should more often play out in favour of the queen than the workers.

Several studies have invoked the disposable soma theory to explain lifespan differences between queens and workers (Jemielity *et al*., 2005; Lucas & Keller, 2014; Kramer *et al*., 2016; da Silva, 2019). The disposable soma theory suggests that senescence or biological deterioration is a consequence of a resource allocation trade-off between resource investment into maintenance vs. reproduction, which can lead to ageing (Kirkwood, 1977). The disposable soma theory might be particularly appealing to explain caste-specific ageing because in social insects, resource allocation trade-offs do not only exist at the within-individual level, but also between individuals of different castes. This could make large resource amounts available to the queen for maintenance and reproduction (Kramer *et al*., 2016), passed on to her by the workers to enhance her reproductive success and thereby their indirect fitness. The few theoretical studies into the evolution of such within-colony resource allocation have modelled how colonies should optimally invest in worker maintenance vs. worker production to derive predictions for worker longevity (Kramer & Schaible, 2013b; Lemanski & Fefferman, 2018). In contrast to the studies mentioned above (Kreider *et al*., 2021; Kramer *et al*., 2022), these theoretical studies confirmed a negative effect of extrinsic mortality on optimal intrinsic worker lifespan. However, these models did not consider the evolution of resource allocation between queen and worker survival. Therefore, they did not explore whether the evolution of between-caste resource allocation can cause caste-specific ageing. In addition, these models did not allow resource allocation to vary with age, although age-specific resource allocation of the queen and workers could crucially impact their survival and lifespan. Hence, there is a need for theoretical predictions on how age-specific resource allocation of queens and workers evolves and if it can lead to caste-specific ageing in social insects.

Several social insect species exhibit an age-related worker division of labour, termed age polyethism, where workers specialise on different tasks throughout their life. Since workers transition from tasks inside the nest towards riskier tasks outside the nest as they get older, extrinsic mortality risk increases with the worker’s age (Seeley, 1982; Tripet & Nonacs, 2004; Rueppell *et al*., 2007; Hartmann *et al*., 2020). It has been shown that the transition from tasks inside the nest to tasks outside the nest is accompanied by a depletion of fat reserves, which could trigger the onset of foraging (Toth *et al*., 2005; Toth & Robinson, 2005; Bernadou *et al*., 2018, 2020). A theoretical model has demonstrated that a division of labour can indeed emerge spontaneously in groups of individuals that are triggered to forage as their nutritional reserves decline, provided that the individuals share resources among each other (Kreider *et al*., 2022).

However, it is unclear whether nutrition-induced foraging and resource sharing can also lead to the emergence of age-related task specialisation. If this is the case, then the resource allocation of queens and workers could be a mechanism for both the emergence of caste-specific ageing and worker age polyethism.

Here, we present an individual-based model for the evolution of ageing in eusocial insects, inspired by the disposable soma theory of ageing. We investigate (1) if caste- or task- and age-specific resource allocation evolves such that lifespan differences between queens and workers emerge; (2) whether the evolution of resource allocation leads to the emergence of worker age polyethism; (3) how extrinsic mortality risk affects the evolution of intrinsic queen and worker lifespans.

## Methods

### General model setup

The evolutionary individual-based model represents a population of *N* eusocial colonies, each inhabited by a queen and workers (parameter settings in Table 1). A key trait in our model is the nutritional state of each queen and worker *x*, which ranges from 0 to 100 arbitrary units (excess resources above the maximum are discarded). The nutritional state of an individual is influenced by the resource allocation decisions by the queen and workers, and it in turn determines the workers’ task choice between foraging or nursing the brood as well as the queen’s and workers’ survival. Apart from a nutritional state, which reflects an individual’s internal state, individuals also have a temporary resource storage organ (e.g. a crop or “social stomach” as in social insects), from which resources can either be digested to improve one’s own nutritional state or shared with other individuals (see Fig. 1F for the different allocation possibilities). This temporary storage organ also has a maximum capacity of 100 arbitrary units. Every time step, each foraging worker collects resources outside the nest, sampled from a normal distribution with mean *μ*_*R*_ and standard deviation *σ*_*R*_, which she invests in increasing her own nutritional state (proportion *p*_1_) or shares with one nursing worker (proportion 1 −*p*_1_). If a foraging worker attempts to transfer resources to a nursing worker whose temporary storage organ is full, she will keep the remainder of resources that the nursing worker cannot receive. Each nursing worker uses the resources she obtained to increase her own nutritional state (proportion *p*_2_), or she gives them away (proportion 1 − *p*_2_) to the brood (proportion (1 −*p*_2_)*p*_3_) or to the queen (proportion (1 − *p*_2_)(1 − *p*_3_)). The queen receives the resources and uses them to increase her nutritional state. She invests a proportion *p*_4_ of her nutritional state into laying eggs, i.e. the resources from the queen’s temporary storage organ first need to undergo “physiological conversion” before they can be invested into egg production. The nutritional state of all individuals declines additively by the metabolic cost of *r* per time step (model predictions for different metabolic costs in Fig. S1).

**Figure 1.**
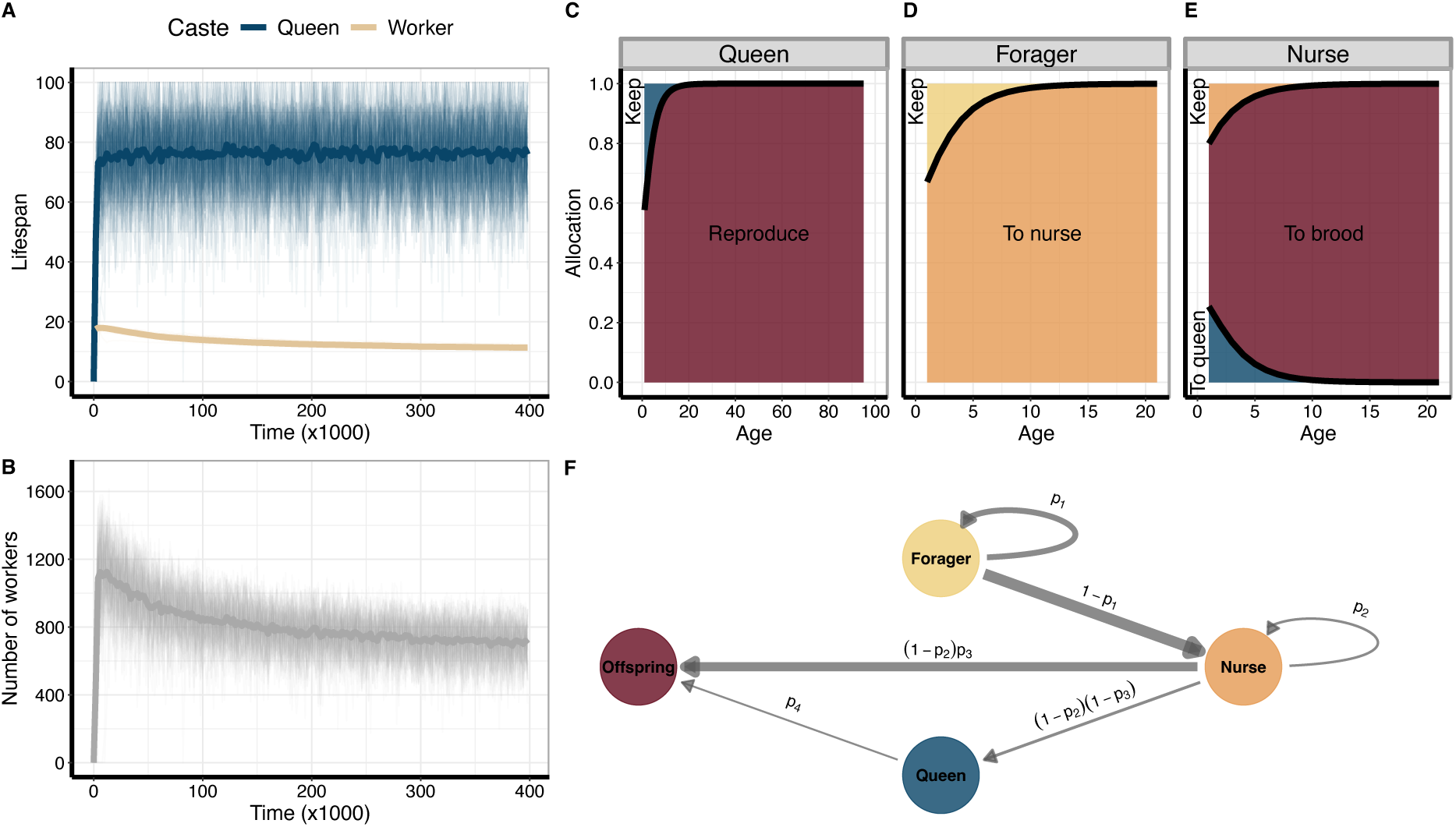
The evolution of resource allocation and caste-specific ageing in eusocial insects. **(A)** Change of intrinsic queen (blue) and worker (sand) lifespans over simulation time. **(B)** Change of number of workers (“colony size”) per colony over simulation time. **(A + B)** The thick line represents the mean over replicate simulations (n = 100). Each thin line is the average at a given time point over all individuals and colonies in one simulation. **(C)** Evolved resource allocation of queens as a function of queen age. The blue area is the resource proportion that the queens keep for themselves, whereas the red area signifies the resource proportion that the queens invest into reproduction (*p*_4_). **(D)** Evolved resource allocation of foragers as a function of their age. The yellow area is the resource proportion that the foragers keep for themselves (*p*_1_), whereas the orange area signifies the resource proportion that they try to pass on to nurses (1 − *p*_1_). **(E)** Evolved resource allocation of nurses as a function of their age. The orange area is the resource proportion that the nurses keep for themselves (*p*_2_). The red area signifies the resource proportion that they pass on to the brood ((1 − *p*_2_)*p*_3_). The blue area is the resource proportion that they pass on to the queen ((1 − *p*_2_)(1 − *p*_3_)). **(C + D + E)** The thick lines represent the mean evolved resource allocation natural cubic spline function averaged over all simulations. The spline functions are cut off at the 95^th^ percentile of queen and worker lifespans, respectively. **(F)** Resource flows within colonies. Arrows indicate how resources can be allocated. Their thickness corresponds to the overall amount of resources transferred from colony founding until queen death, averaged over simulations. Note that the queen’s investment into offspring occurs by production of eggs whereas the nurses’ investment in offspring occurs by directly feeding them.

**Table 1.** Parameter values used in the simulations, unless stated otherwise.

| Parameter | Value | Meaning |
| --- | --- | --- |
| $N$ | 1000 | Number of colonies |
| $\mu_R$ | 70.0 | Mean resources obtained per foraging trip |
| $\sigma_R$ | 5.0 | Standard deviation of resources obtained per foraging trip |
| $r$ | 10.0 | Metabolic cost, i.e. decline of nutritional state per time step |
| $m$ | 0.03 | Mutation rate |
| $\sigma_m$ | 0.01 | Standard deviation for mutational effect |
| $s_f$ | 0.1 | Scaling parameter for steepness of function $f$ |
| $l_f$ | 50.0 | Scaling parameter for location of function $f$ |
| $c$ | 1.0 | Egg production cost |
| $\mu_a$ | 60 | Mean nutrition at which larvae emerge as adults |
| $\sigma_a$ | 5.0 | Standard deviation nutritional state at which larvae emerge |
| $s_d$ | 0.1 | Scaling parameter for steepness of function $d_{\text{int}}$ |
| $l_d$ | 10.0 | Scaling parameter for location of function $d_{\text{int}}$ |
| $a_{\text{max}}$ | 100 | Maximum age |
| $d_{\text{ext}}$ | 0.1 | Probability of extrinsic mortality during foraging |
| $q$ | 10 | Foraging trips of the queen in the colony founding phase |

### Evolving traits

The resource investment of queens, foraging workers and nursing workers is age-specific, e.g. the proportions *p*_1_to *p*_4_ are each a function of the age of the individual. We model these age-specific expressions using highly flexible natural cubic spline functions (details in supplementary materials) (Lagos-Oviedo *et al*., 2024; Rees-Baylis *et al*., 2024). Their shape is determined by the coefficients or weights of a fixed number of basis functions. We deploy five such basis functions, whose coefficients are genetically determined by five pairs of homologous genes. Each gene value mutates with probability *m* upon every meiotic event. Mutational effects are drawn from a normal distribution with a mean of zero and a standard deviation of *σ*_*m*_. We initialise the gene values such that resource allocation is initially equal across all allocation possibilities and identical across ages. We assume haplodiploid genetics, where females are diploid and males are haploid. We assume additive gene action. Full recombination occurs during the production of gametes, assuming no linkage between the genes.

### Worker task allocation

We adapted the division of labour model by Kreider *et al*. (2022), where a sufficiently low nutritional state triggers individuals to forage. The probability, that a worker forages, is given by the logistic function

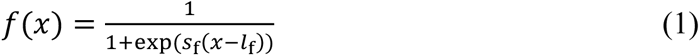

where *S_f_* is a scaling parameter for the steepness of the function and *l*_f_ determines the function’s location, i.e. the nutritional state at which individuals forage with a probability of 0.5.

### Reproduction

Queens use the resources they invest in reproduction to lay eggs. Each egg has a production cost of *c*. As many offspring are fed to maturity as the amount of resources allocated to the brood allows. Offspring mature to adulthood at a nutritional state sampled from a normal distribution with mean *μ*_a_ and standard deviation σ_a_. The remaining offspring, that did not receive sufficient resources, die.

### Survival

Queens and workers die when their nutritional state gets too low. The intrinsic probability of death per timestep *d*_int_is given by

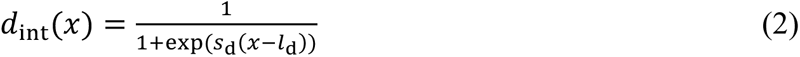

where *S*_d_is a scaling parameter for the steepness of the function and *l*_d_ determines the function’s location, i.e. the nutritional state at which individuals die with a probability of 0.5. The age of individuals is restricted to *a*_max_. In addition, foraging workers die from extrinsic mortality risk with probability *d*_ext_. Thus, the total probability of death of a foraging worker is *d*_int_ + (1 − *d*_int_)*d*_ext_.

### Colony life cycle

Each colony is founded by a single queen that forages *q* times to provision her first offspring with resources. After the emergence of workers, the queen stays in the nest, whereas her workers forage or nurse the brood. The queen resumes foraging and provisioning brood if all workers die. If a queen dies, the workers also die. A new founding queen is sampled from the provisioned larvae in the surviving colonies. The founding queen mates with one male, which is mothered by another queen sampled proportionally to the number of provisioned larvae in the colonies.

### Model analysis

We simulated the model for 400 000 timesteps. We recorded average intrinsic lifespans of queens and workers, who died because their nutritional state became low (eq. 2) and excluding deaths from extrinsic mortality, and number of workers per colony every 100 timesteps. At the end of each simulation, we obtained data from 50 colonies on the evolved allocation strategies (age-specific gene values of *p*_1_to *p*_4_), the overall resource flows between all allocation possibilities (Fig. 1F) over the lifespan of the queen in her colony, and the nutritional state and task choice at each age of every individual in the colony. We calculated the queen/worker lifespan ratio on mean intrinsic lifespans of each simulation. We analysed simulation results in R 4.5.2 (R Core Team, 2025) using the packages *tidyverse* (Wickham *et al*., 2019), *cowplot* (Wilke, 2025), *igraph* (Csárdi & Nepusz, 2006; Antonov *et al*., 2023; Csárdi *et al*., 2026), *ggraph* (Pedersen, 2025), and *MetBrewer* (Mills, 2022).

## Results

### Caste-specific ageing

Lifespan differences between queens and workers evolve readily in the simulations and the number of workers in colonies increases initially (Fig. 1A+B). After an initial increase, worker lifespans decrease slowly, also causing a slow decrease in the number of workers in colonies. At the end of the simulations, the median queen/worker lifespan ratio was 6.76 [95% CI: 6.53–6.98]. The caste differences in lifespan are a consequence of the evolved resource allocation strategies by the queen and workers. Queens evolve to keep some resources early in their life but evolve to fully invest in reproduction as they get older (Fig. 1C). Foragers keep a small proportion of resources for themselves when they are young but transfer most of their resources to nurses (Fig. 1D). Nurses also keep a small proportion of resources for themselves when they are young and transfer some resources to the queen, while the largest proportion of resources is fed to the offspring (Fig. 1E). The evolved resource allocation results in colony-level resource flows where most resources, handled by the foragers, are transferred to nurses, who in turn transfer most resources to the offspring (Fig. 1F). The amount of resources transferred to the queen seems relatively small. However, Fig. 2A shows that this resource amount is sufficient to fully replenish the queen’s nutritional state after an initial colony founding period, where her nutritional state is lower because of resource limitation due to a smaller number of workers (queen age coincides with time since colony founding).

**Figure 2.**
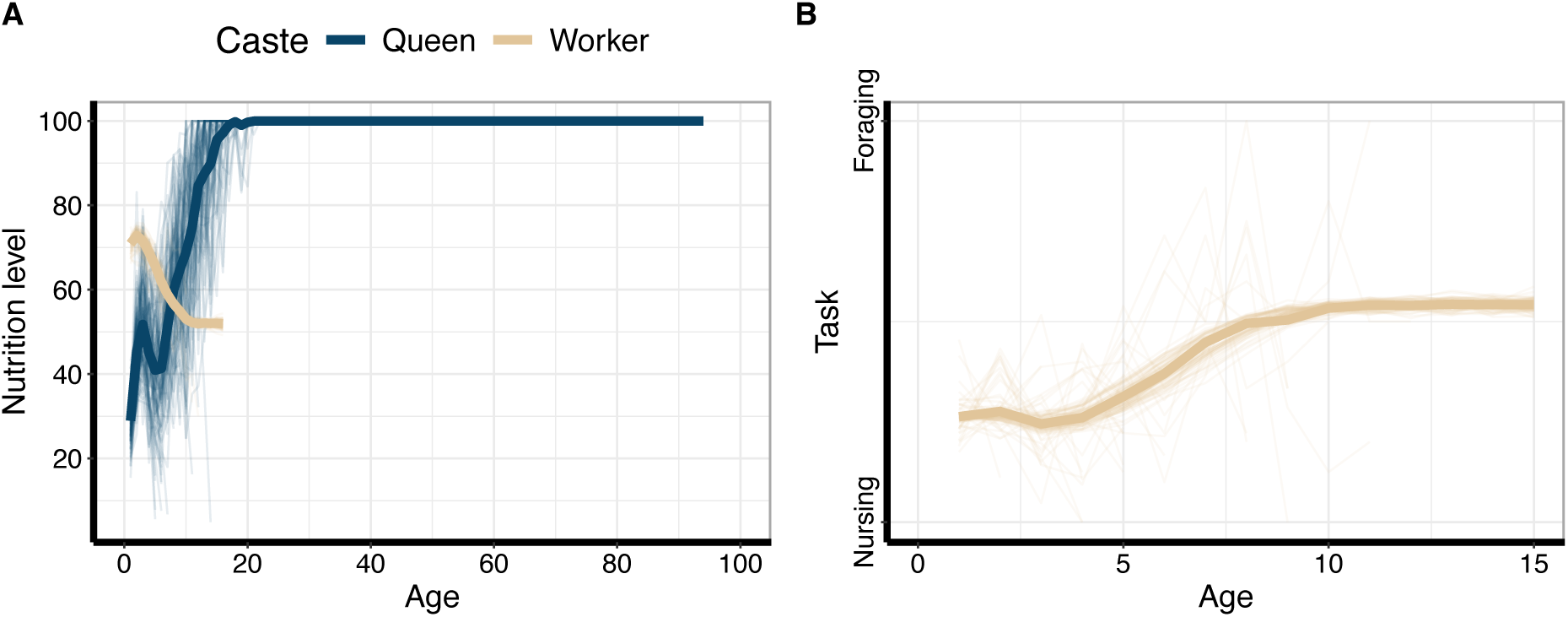
Dynamics of nutritional state with age and age-related worker division of labour. **(A)** Nutritional state dynamics by queen (blue) and worker (sand) age. Note that queens have a lower nutrition level at age 0 than workers because queens immediately invest in offspring production. **(B)** Worker task choice between nursing and foraging varies with age. **(A + B)** The thick line represents the mean over simulations. The thin lines are the means from individual simulation replicates. All lines are cut off at the 95^th^ percentile of queen and worker lifespans, respectively.

### Age polyethism

The nutritional state of workers declines as they get older (Fig. 2A). Young workers are more likely to nurse the brood and become more likely to forage when older (Fig. 2B). Consequently, an age-related worker division of labour, i.e., worker polyethism, emerges. An assumption in our parameter choice is that workers emerge with a high nutritional state, which is realistic (Toth *et al*., 2005; Bernadou *et al*., 2018, 2020). This predisposes young workers to nurse. We also investigated a model scenario where workers emerge with a low nutritional state, which predisposes them to forage early in life. However, even in this scenario, workers follow a similar age-related division of labour as in Figure 2, after they have foraged and increased their nutritional state when they are young (Fig. S2).

### Extrinsic mortality

Lifespan differences between queens and workers evolve in the absence of extrinsic mortality risk (Fig. 3). Increasing extrinsic mortality risk causes a decline in intrinsic worker lifespans but no change in intrinsic queen lifespans.

**Figure 3.**
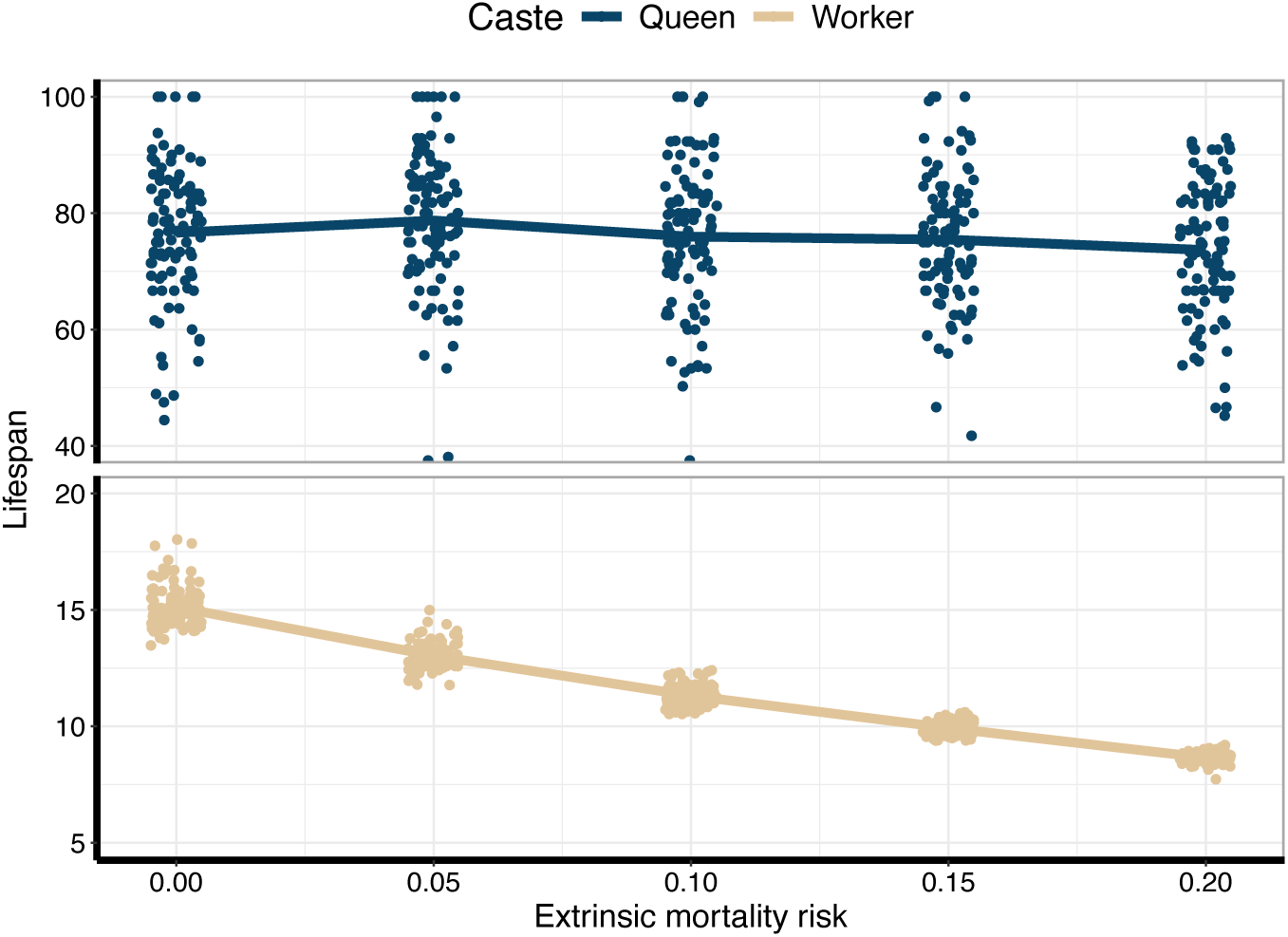
Effect of extrinsic mortality risk on the evolution of caste-specific ageing. Evolved intrinsic queen (blue) and worker (sand) lifespans (note the different y-axes scales for queens and workers for comparability of queen lifespans and worker lifespans between simulations with different extrinsic mortality risk). The line connects the means over replicate simulations (n = 100). Each dot represents the mean lifespan from one simulation. The dots are jittered horizontally.

## Discussion

The disposable soma theory of ageing has often been invoked to explain the evolution of divergent lifespans between queens and workers of eusocial insects (Jemielity *et al*., 2005; Lucas & Keller, 2014; Kramer *et al*., 2016; da Silva, 2019). However, to date no theoretical model has investigated whether resource allocation dynamics within colonies evolve such that differences in queen and worker lifespans emerge as a result. Here, we demonstrate that evolved resource allocation strategies by queens and workers lead to an emergence of long-lived queens and short-lived workers. Our model predicts a median queen/worker lifespan ratio of 6.76 [95% CI: 6.53–6.98]. This prediction matches lifespan estimates for monogynous eusocial insects, where the median queen/worker lifespan ratio is 5.0 [95% CI: 4.0–7.8] (Kramer & Schaible, 2013a; Kramer *et al*., 2022). In addition to predictions for caste-specific ageing, we also demonstrate that the evolved resource allocation dynamics can lead to an age-related division of labour (age polyethism). Our model therefore demonstrates that caste-specific ageing and age-related division of labour can emerge from a shared mechanism, which is the evolved within-colony resource allocation.

Trade-offs between survival and fecundity have often been assumed to be virtually absent in eusocial insects, because queens are both long-lived and highly fertile, whereas workers are short-lived and often sterile (Kramer *et al*., 2015; Korb, 2016; Lin *et al*., 2021; Zug *et al*., 2025). Even comparisons between queens from different colonies have demonstrated a positive correlation between survival and fecundity (Lopez-Vaamonde *et al*., 2009; Heinze *et al*., 2013; Schrempf *et al*., 2017; Zug *et al*., 2025). Consequently, the trade-off between survival and fecundity has often been assumed to be “reversed” in eusocial insects (Parker, 2010; Séguret *et al*., 2016; Korb *et al*., 2021; Korb & Heinze, 2021; Rau *et al*., 2023). However, such a positive correlation does not imply the absence of a trade-off between fecundity and survival (Van Noordwijk & De Jong, 1986; Zug *et al*., 2025). Instead, the trade-off is masked because stochasticity causes some colonies to be more successful which lets them accumulate more resources that are available for reproduction and queen survival (Zug *et al*., 2025). In our model, the trade-off between survival and fecundity is a crucial ingredient – the same resource cannot be simultaneously invested in survival and fecundity. Since sterile workers can only enhance their inclusive fitness through enhancing the queen’s reproductive output, the workers evolve to pass on resources to the queen and the offspring, which has a negative effect on their survival. Previous resource-based models for the evolution of social insect ageing have predicted optimal worker lifespans under different rates of extrinsic mortality risk but did not explicitly model allocation effects on queen lifespans (Kramer & Schaible, 2013b; Lemanski & Fefferman, 2018). Our resource-based model is the first to demonstrate that between-caste trade-offs can lead to the evolution of caste-specific ageing. Consequently, the trade-off between survival and reproduction is not absent in eusocial insects, but it can be crucial for the evolution of caste-specific ageing.

Division of labour modelling has shown that resource sharing can lead to the spontaneous emergence of division of labour, when foraging is triggered by a low nutritional state (Kreider *et al*., 2022). This is because foraging individuals are more likely to be triggered to forage again if they give resources away, whereas nursing individuals delay their onset of foraging if they receive resources. We here extended this model by letting the proportion of resources shared evolve and by allowing for task-specific extrinsic worker mortality. Our model shows a decrease of worker nutritional state with age, as found in eusocial insects (Bernadou *et al*., 2020), and demonstrates the emergence of an age-related division of labour, which is widespread across eusocial insects (Seeley, 1982; Rueppell *et al*., 2007; Hartmann *et al*., 2020).

It has long been assumed that the evolution of caste-specific ageing is caused by differential exposure to extrinsic mortality risk of queens and workers (Keller & Genoud, 1997). However, theoretical modelling of the evolution of ageing in social insects has shown that caste-specific ageing evolves due to reproductive division of labour and irrespective of caste-specificity of extrinsic mortality risk (Kreider *et al*., 2021; Kramer *et al*., 2022). Our model supports these findings since differences in queen and worker lifespans also evolve in the absence of extrinsic mortality. However, in contrast to previous models which assumed genes with direct age-dependent effects on survival (Kreider *et al*., 2021; Kramer *et al*., 2022), resource-based ageing models (Kramer & Schaible, 2013b; Lemanski & Fefferman, 2018), including the model presented here, do find a decrease in worker lifespan with increasing extrinsic mortality risk, even in monogynous colonies with sterile workers. This raises the question why different types of models yield different predictions. An increase in extrinsic mortality risk should cause workers to evolve to give more resources away, since resources used to increase their own nutritional state would be wasted if the workers died from extrinsic mortality. This effect is likely reinforced in our model because workers are more likely to forage as they get older. Hence, exposure to extrinsic mortality risk is “concentrated” in particular worker age classes, causing workers to evolve to pass on more resources (Fig. S3), which also leads to a higher probability of intrinsic death. Evolving to pass on resources to other individuals in the colony in anticipation of one’s own death to enhance one’s own inclusive fitness can be viewed as adaptive ageing (Galimov & Gems, 2021), where a higher level of intrinsic mortality is adaptive, since it makes resources available for the queen’s reproductive output.

An analogy between eusocial insect colonies and multicellular organisms is frequently made, where the reproductive queen is assumed to be analogous to the germline and the workers analogous to the somatic cells (Helanterä, 2016; Boomsma & Gawne, 2018). This analogy is also useful in the context of the evolution of ageing (Bernadou *et al*., 2021; Kramer *et al*., 2022). Queens are extraordinarily long-lived, analogously to the “immortal” germline (as classically assumed; Weismann, 1892), whereas the workers are short-lived and similarly disposable as the soma. In multicellular organisms, apoptosis (programmed cell death) has evolved to eliminate senescent or damaged cells and prevent them from cancerous proliferation (Fernald & Kurokawa, 2013). Akin to the adaptively evolved resource transfers by old foragers to younger individuals, which lead to a higher level of intrinsic mortality in our model, apoptotic cells release metabolites that can be used by neighbouring cells influencing their states (Medina *et al*., 2020). The co-dependency of castes in superorganismal eusocial insects and cell types within multicellular organisms has likely similarly led to the evolution of mechanisms that increase intrinsic mortality, that is adaptive ageing, in specific castes or cell types to enhance the fitness of the (super-)organism.

In summary, our model extends the disposable soma theory of ageing to eusocial insects and demonstrates that the evolution of resource allocation between queens and workers leads to caste-specific ageing. Simultaneously, the evolution of resource allocation between foraging and nursing workers also leads to an age-related division of labour. Hence, our model demonstrates that caste-specific ageing and age polyethism emerge readily from a shared mechanism: the evolution of resource allocation.

## Author contributions

All authors contributed to conceptualising this research. TJ and JJK implemented the model. JJK and BHK analysed the model. JJK wrote the manuscript with input from all authors.

## Acknowledgements

We would like to thank the Center for Information Technology of the University of Groningen for providing access to the Hábrók high performance computing cluster. JJK was supported by an EMBO postdoctoral fellowship (ALTF 786-2024).

