## Supplement for "Caste-specific ageing emerges from the evolution of resource allocation in eusocial insects"

### Evolving natural cubic splines

We model resource allocation as a function of age $a$ using flexible smooth natural cubic spline functions (Lagos-Oviedo *et al.*, 2024; Rees-Baylis *et al.*, 2024). These functions, which consist of connected cubic polynomials that are linear in their tail ends, follow the definition by Harrell (2001). A natural cubic spline function has $k$ knots ($k = 5$), which are the connection locations of the cubic polynomials with respect to $a$. We assume that these locations are evenly distributed across $a$. The knot locations of the $k$ knots are $t_1, \dots, t_k$. The natural cubic spline function, consisting of the basis functions $B_i$, is then given by

$$f(a) = \beta_0 B_0 + \beta_1 B_1 + \beta_2 B_2 + \cdots + \beta_{k-1} B_{k-1}, \tag{S1}$$

where $B_0 = 1$ and $B_1 = a$. The remaining terms $B_2, \dots, B_{k-1}$ are calculated by iterating over $j = 1, \dots, k-2$ according to

$$B_{j+1} = \left(a - t_j\right)_+^3 - \frac{(a-t_{k-1})_+^3 (t_k - t_j)}{t_k - t_{k-1}} + \frac{(a-t_k)_+^3 (t_{k-1} - t_j)}{t_k - t_{k-1}}, \tag{S2}$$

where $(\dots)_+$ indicates that a term is set to 0 when it is evaluates to a number below 0. The natural cubic spline function can also be written in matrix notation

$$f(a) = \begin{bmatrix} B_0(a_1) & B_1(a_1) & B_2(a_1) & \cdots & B_{k-1}(a_1) \\ B_0(a_2) & B_1(a_2) & B_2(a_2) & \cdots & B_{k-1}(a_2) \\ \vdots & \vdots & \vdots & \ddots & \vdots \\ B_0(a_n) & B_1(a_n) & B_2(a_n) & \cdots & B_{k-1}(a_n) \end{bmatrix}_a \begin{bmatrix} \beta_0 \\ \beta_1 \\ \beta_2 \\ \vdots \\ \beta_{k-1} \end{bmatrix} \tag{S3}$$

Here, the matrix is the basis matrix of the natural cubic spline function. The column vector $\beta$ contains the parameters $\beta_0, \dots, \beta_{k-1}$ for the natural cubic spline function. The basis matrix is precalculated for every age $a$. The natural cubic spline function parameters in column vector $\beta$ are the evolving gene values. To evaluate the phenotypic value at age $a$, we multiply the column vector $\beta$ with the $a$-th row of the basis matrix (as signified by basis matrix subscript $a$). This value is subsequently logistically transformed to result in a proportion. We initialise all values of $\beta = 0.0$, which results in equal allocation to oneself vs. someone else across ages. The only exception to this is the allocation of nurses, which have three allocation options (self, brood, queen). Here, $\beta_0 = -ln\ (2)$, which after logistic transformation results in $0.\overline{3}$.

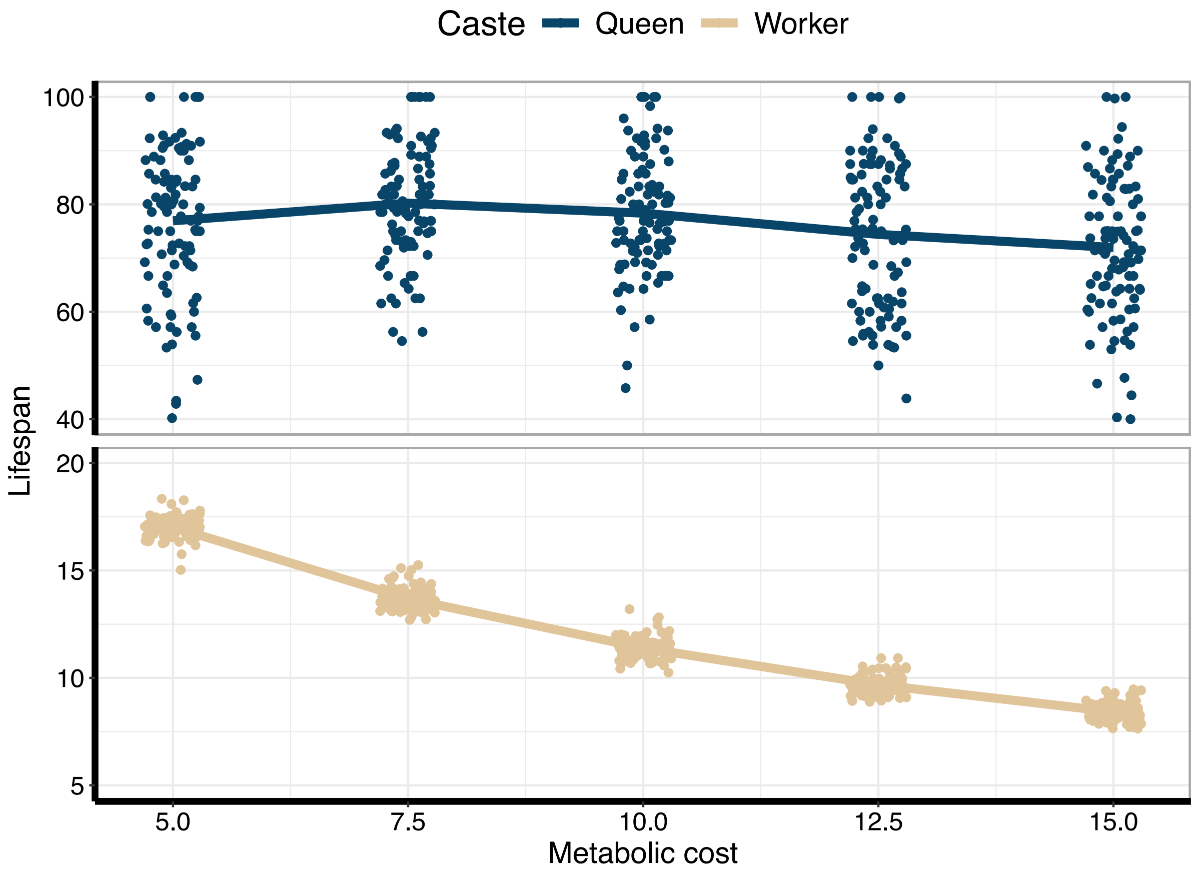


**Figure S1.** Effect of metabolic cost on the evolution of caste-specific ageing. Evolved intrinsic queen (blue) and worker (sand) lifespans (note the different y-axes scales for queens and workers). The line connects the means over replicate simulations (n = 100). Each dot represents the mean lifespan from one simulation. The dots are jittered horizontally.

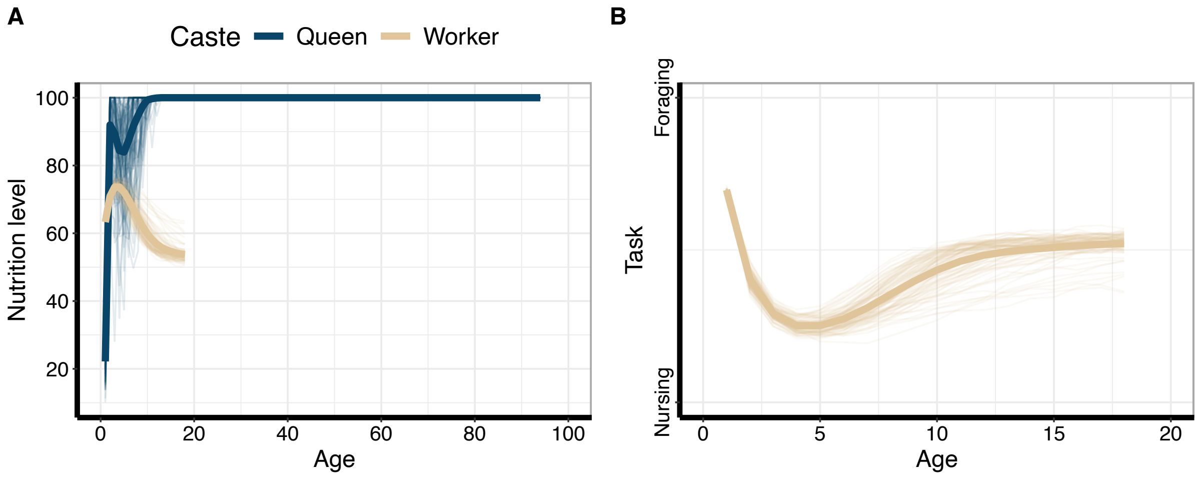


**Figure S2.** Nutrition levels over age and age-related worker division of labour, when workers emerge with a low nutrition level. **(A)** Change of nutrition level with queen (blue) and worker (sand) age. **(B)** Worker task choice between nursing and foraging depends on age. **(A + B)** The thick line represents the mean over simulations. The thin lines are the means from individual simulations. All lines are cut off at the 95th percentile of queen and worker lifespans, respectively.

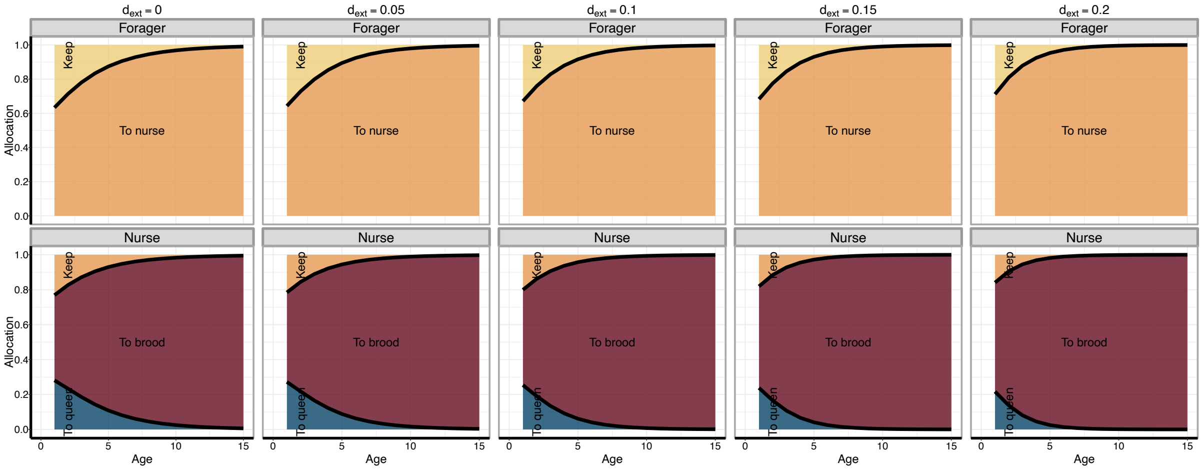


**Figure S3.** Evolved age-specific resource allocation of foragers and nurses under different levels of extrinsic mortality risk ($d_{\text{ext}}$). The yellow area is the resource proportion that the foragers keep for themselves ($p_1$), whereas the orange area in the upper row signifies the resource proportion that they try to pass on to nurses ($1 - p_1$). The orange area in the lower row is the resource proportion that the nurses keep for themselves ($p_2$). The red area signifies the resource proportion that they pass on to the brood ($(1 - p_2)p_3$). The blue area is the resource proportion that they pass on to the queen ($(1 - p_2)(1 - p_3)$). The thick lines represent the mean evolved resource allocation natural cubic spline function averaged over all simulations (n = 100 per parameter setting). The x-axis is cut-off at an age of 15 to better show the decrease of the proportion of resources kept by workers with increasing extrinsic mortality risk.
